# Targeting Oncogenic Mutations in Colorectal Cancer using Cryptotanshinone

**DOI:** 10.1101/2020.09.15.298398

**Authors:** Haswanth Vundavilli, Aniruddha Datta, Chao Sima, Jianping Hua, Rosana Lopes, Michael Bittner

## Abstract

Colorectal cancer (CRC) is one of the most prevalent types of cancer in the world and ranks second in cancer deaths in the US. Despite the recent improvements in screening and treatment, the number of deaths associated with CRC is still very significant. The complexities involved in CRC therapy stem from multiple oncogenic mutations and crosstalk between abnormal pathways. This calls for using advanced molecular genetics to understand the underlying pathway interactions responsible for this cancer. In this paper, we construct the CRC pathway from the literature and using an existing public dataset on healthy vs tumor colon cells, we identify the genes/pathways that are mutated and are possibly responsible for the disease progression. We then introduce drugs in the CRC pathway, and using a boolean modeling technique, we deduce the drug combinations that produce maximum cell death. Our theoretical simulations demonstrate the effectiveness of Cryptotanshinone, a traditional Chinese herb derivative, achieved by targeting critical oncogenic mutations and enhancing cell death. Finally, we validate our theoretical results using wet lab experiments on HT29 and HCT116 human colorectal carcinoma cell lines.

## 1 Introduction

Colorectal cancer (CRC) is the third most common type of cancer worldwide and its incidence is alarmingly increasing in individuals less than 50. The American Cancer Society estimates that in 2020 alone, the US will have around 148000 cases and 53000 deaths attributed to CRC [1]. Colorectal cancer was the most prevalent cause of cancer death in the US around the 1950s, but the numbers have since come down, in part because of early detection, better treatment, and the adoption of healthy lifestyles. Although overall CRC cases and mortality rates continue to decline, this progress is increasingly restricted to older age groups [2]. Hence, it is important to design efficient and cutting-edge techniques for fighting colorectal cancer.

Cancer is a type of disease that develops and grows when a living cell’s control system becomes faulty. Old cells, which are supposed to undergo programmed cell death, do not die and grow out of control and eventually form a tumor [3]. These tumorous cells that can invade other tissue, and subsequently acquire the capability to metastasize and spread to other parts of the body, constitute malignant tumors. Most patients with colorectal cancer die because of malignant tumors and the cancer spreading to other organs. This highlights the need for a better and comprehensive understanding of the underlying oncogenic mutations and interactions responsible for the progression of this disease.

Multicellular organisms have highly advanced communication networks to coordinate various biological processes. Among them, gene regulatory networks (GRN), which characterize the interactions between genes and other molecules, play an important role in managing biological processes such as cell proliferation, metabolism, cell differentiation, and apoptosis [4]. In a normal cell, these processes are tightly regulated, and misregulations in them (also called mutations) have been linked to diseases such as cancer [5].

Surgery and chemotherapy are the two commonly used methods for treating cancer [6]. Although surgery is the most frequently used method, unfortunately, it has limitations when the tumor is inoperable due to its size or if metastases have occurred making it infeasible to surgically operate on all the tumors. Chemotherapy, on the other hand, makes use of external cytotoxic agents like therapeutic drugs or radiation to destroy the tumor [7]. Therapeutic drugs are designed to intervene at specific locations by binding to explicit target(s). However, since the number of possible mutations is vast, there is a demand for finding a few “significant” genes that can be controlled rather than a large group of genes. The motivation for this is that designing experiments and drugs to control a large number of genes is both costly and tedious when compared to the effort involved in doing the same for a few crucial genes. However, since finding these crucial genes is not straightforward, scientists have been developing mathematical models and techniques to facilitate their identification. The key goal of these modeling techniques in cancer is to provide a basis for designing chemotherapy regimes and to facilitate qualitative prognosis about the dynamic evolution of the disease. Various techniques such as differential equations [8], boolean networks [9]–[11], probabilistic boolean networks [12], and bayesian networks [13]–[15] have had some success along these lines.

In this paper, we use bayesian and boolean approaches respectively to arrive at the aberrant pathways and the best drug combinations to target colorectal cancer. Using a public dataset of healthy versus tumor colon cells, we first analyze the underlying genes and pathways that are possibly responsible for CRC. We then theoretically calculate the effectiveness of different drugs and their combinations by simulating a boolean network model of CRC and ranking the drugs based on their performance to accomplish cell death.

The paper is organized as follows. In section 2, we discuss the dataset and methodology used. Then, in section 3, we build and analyze the colorectal cancer pathway from the biological literature. In section 4, we present the simulations, theoretical results, and experimental validation on CRC cell lines. Finally, in section 5, we discuss the biological relevance of our results and provide some concluding remarks.

## 2 Methodology

### Dataset

We used an existing public dataset GSE44076 available in NCBI’s Gene Expression Omnibus [16]. The dataset contains 50 healthy colon samples and 98 tumor samples. The samples of healthy subjects were obtained without colonic lesions at the time of colonoscopy and the tumor samples were obtained from subjects with histologically confirmed diagnosis of colorectal adenocarcinoma.

We are interested in analyzing how the gene expression values in healthy samples compare against the values in the tumor samples.

### Probabilities

Several studies show that quantizing the gene expression values into a binary format of 0s (inactive) and 1s (active) offers advantages like computational lucidity and robustness against noise. Hence, for each gene in the colorectal cancer pathway, we first discretized its gene expression values using a binary framework. We used the maximum likelihood estimator for the mean *µ* as the threshold. This threshold is justified by the law of large numbers and the central limit theorem.

Then, we used a bayesian framework of prior and posterior probabilities to estimate the network parameters [17]. We chose a non-informative uniform beta distributed prior and since the beta distribution is conjugate to the binomial likelihood, the conditional posterior probability distributions are also beta distributed. Hence, the probability that a gene is over-expressed is given by the ratio of over-expressed samples to the total number of samples in that dataset [18]. That is,

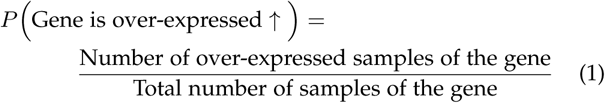

For every gene in the CRC pathway, we computed this probability separately for healthy and tumor samples from the GSE44076 dataset and evaluated the ratio. We define the ratio of probabilities (ROP) as,

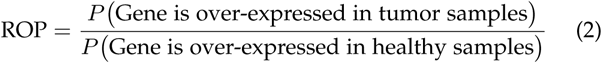

Clearly, a ratio greater than 1 (i.e. ROP *>* 1) suggests that a particular gene is operating in an aberrantly up-regulated fashion in the tumor samples as compared to the healthy samples. Similarly, a ratio less than 1 (i.e. ROP *<* 1) suggests that a particular gene is operating in an aberrantly down-regulated fashion in the tumor samples as compared to the healthy samples.

### Boolean modeling

Boolean network modeling is a common technique to design discrete-time, discrete-space biological systems including gene regulatory networks (GRN). Boolean networks represent a paradigm that can be used to integrate pathway information to model the complete dynamic behavior of the cell [19]. Each node in a boolean network can be in one of two binary states (inactive or active). We assign a a ‘1’ value to the active state and a ‘0’ value to the inactive state.

In a GRN, the activity of a gene is determined by one or more genes. These interactions among multiple genes can be modeled as boolean logic functions, where the nodes represent the genes and the edges represent the interactions among the genes. For example, if a gene is independently activated by two genes, that interaction can be modeled using an OR gate.

We now construct the colorectal cancer pathway from the literature and discuss the key genes and pathways involved in it.

## 3 Colorectal cancer Pathway

Colorectal cancer results from an accumulation of mutations in oncogenes and tumor suppressor genes. Mutations in the MAPK/ERK pathway, the JAK/STAT pathway, the PI3K/AKT pathway and several other genes/pathways drive tumor progression and metastasis [20]. We shall now look at these underlying processes and malignancies in CRC.

### MAPK/ERK

Mitogen-activated protein kinases (MAPK) pathway is crucial in supervising key cellular functions such as cell growth, survival, and apoptosis [21]. This pathway is activated when epidermal growth factor (EGF) binds to epidermal growth factor receptor (EGFR). Upon this ligand binding, this complex activates growth factor receptor-bound protein 2 (GRB2) in presence of son of sevenless (SOS), which further stimulates RAS, a well known proto-oncogene. Active RAS recruits RAF protein that activates MEK, which in turn activates extracellular regulated kinase (ERK) by phosphorylation [22]. ERK then permeates into the cell nucleus and triggers multiple transcription factors such as FOS and JUN.

In a healthy cell, these processes are overseen in a controlled fashion, but mutations in RAS, EGFR, ERBB2, and other receptor tyrosine kinases have shown increased signaling and overdrive this pathway.

### PI3K/AKT

Another important and complex pathway which involves RAS is the phosphoinositide 3-kinase (PI3K) pathway [23]. The regulatory subunit of PI3K, p85, disinhibits the catalytic subunit, p110, and this phosphorylates PIP2 to PIP3. RAS can also stimulate this pathway by directly binding to p110. Conversely, phosphatase and tensin homolog (PTEN), a tumour suppressor protein, dephosphorylates PIP3 back to PIP2, thus terminating signaling [24]. PIP3 recruits and phosphorylates AKT, that further activates genes in charge of cellular growth and apoptosis.

Deletions in the PTEN gene make it unable to convert PIP3 back to PIP2, and this is responsible for Cowden syndrome, which is an autosomal dominant disorder that influences multiple cancers. Increased PI3K activity and mutations in PTEN have been shown in at least 20%-25% of CRCs [25].

### JAK/STAT

Janus kinase (JAK)/signal transducer and activator of transcription (STAT) signaling pathway is essential for various physiological processes, including immune function, cell survival, and cell growth. STATs transduce signals from cytokines and RTKs such as EGFR [26].

Abnormalities in the JAK/STAT3 pathway with increased activity of STAT3 has been linked to oncogenesis of multiple cancers including colorectal cancer. This decisive role of STAT3 in tumorigenesis and cancer progression makes it an attractive therapeutic target.

### WNT

Wnt signaling supervises the level of the key modulator-catenin for signal transduction through processes comprising ubiquitin-mediated degradation and phosphorylation [27]. This is coordinated by the *β*-catenin complex, which consists of important proteins including glycogen synthase kinase 3 (GSK3), adenomatous polyposis coli (APC), and AXIN. The protein dishevelled (DVL) transduces Wnt signals from receptors to downstream effectors. In the nucleus, *β*-catenin forms an active transcriptional complex with T-cell factor (TCF)/lymphoid enhancer-binding factor (LEF) transcription factors, further activating downstream Wnt target genes [28].

The family of Wnt proteins plays a crucial role in the embryonic development process, the determination of cell fate, proliferation and differentiation of stem cells including human mesenchymal stem cells. Under orderly circumstances, *β*-catenin levels in the cytoplasm are controlled by APC and GSK3. Aberrant gene expression levels of APC and *β*-catenin have been shown to be present in almost all human colon tumors.

### TGF-*β*

The Transforming Growth Factor Beta (TGFB) signaling pathway is one of the most commonly mutated cellular signaling networks in multiple human cancers [29]. TGFB ligands bind to type TGFB receptors which then phosphorylate downstream transcription factors, SMAD2 and SMAD3, allowing them to bind to SMAD4. Activated SMAD complexes translocate into the nucleus and interact with other transcription factors. In addition, TGFB can activate the JNK pathway through MKK4 and drive the proliferation of CRC cells.

There is growing evidence of the role of TGFB in colorectal carcinogenesis as TGFB acts as a tumor promoter and is generally highly expressed. Signaling alterations mediated by mutations in SMADs also contribute to CRC development and progression [30].

### Therapeutic Drugs

Chemotherapeutic drugs are designed to impede cell growth and other processes by altering the functioning of vital genes and proteins. They carry this out by either blocking the function of the proteins they intervene with or by inducing their effect via binding to the target receptor site(s). In table 1, we tabulate the therapeutic drugs that are well known to intervene and bind specific genes in the colorectal cancer pathway along with the relevant supporting literature. In this paper, we will be using these drugs and their combinations in our theoretical simulations and experiments.

**TABLE 1:** Drugs and their gene intervention points.

| Drug | Gene(s) targeted | References |
| --- | --- | --- |
| AG1024 | IGF1R | [31], [32] |
| U0126 | MEK1 | [33] |
| NT157 | IRS1 | [34], [35] |
| LY294002 | PIK3CA | [36] |
| HO-3867 | STAT3 | [37] |
| Lapatinib | EGFR + ERBB2 | [38], [39] |
| Temsirolimus | mTOR | [40] |
| Cryptotanshinone | STAT3 + ERK | [41], [42] |

Using the information discussed above and other literary sources, we constructed the gene regulatory network of colorectal cancer as shown in Fig. 1. The gene interactions are represented by arrows, where a black arrow denotes activation and a red arrow denotes inhibition. We also constructed the boolean equivalent of this CRC pathway as shown in Fig. 2.

**Fig. 1:**
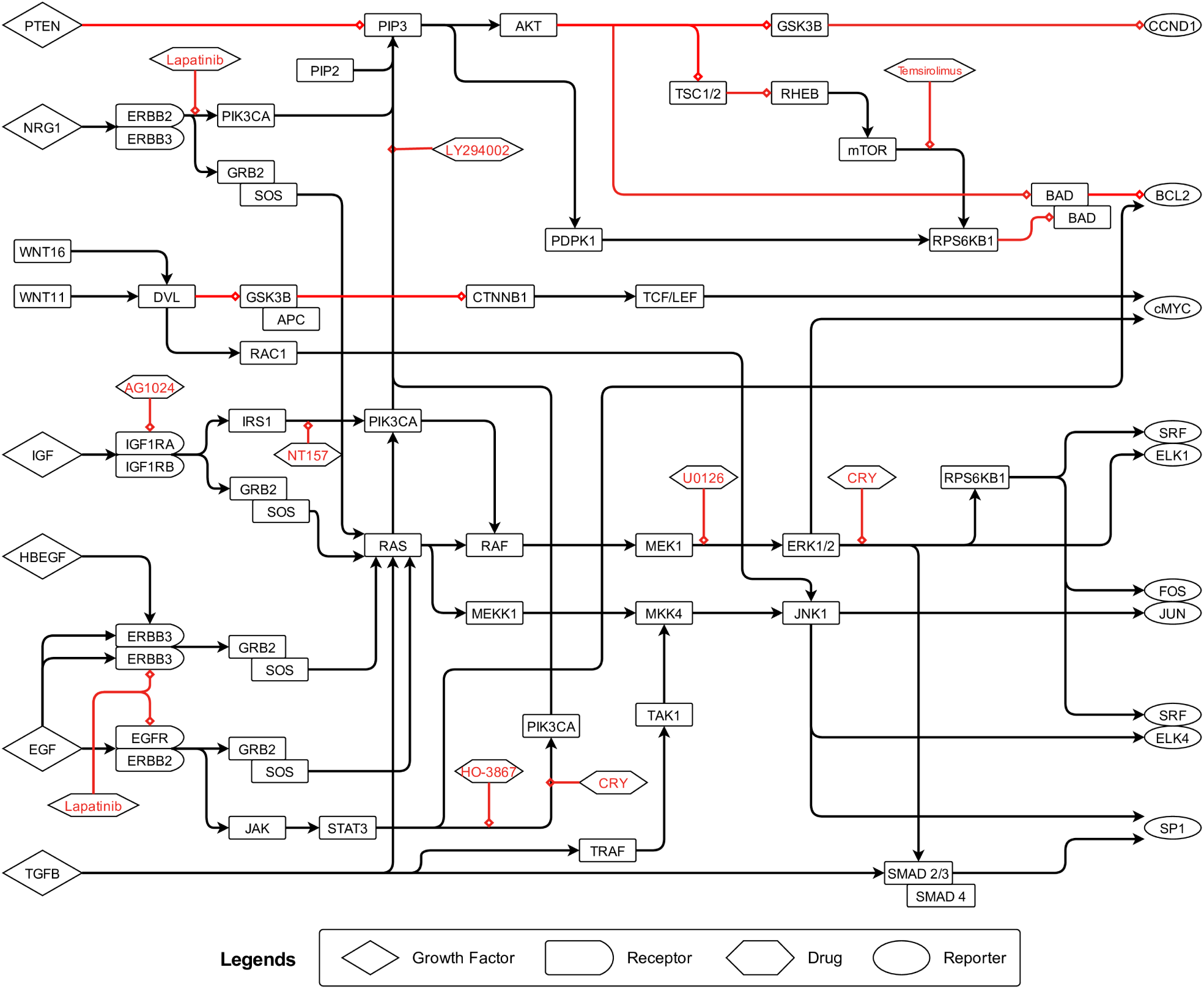
Colorectal cancer pathway. A black arrow denotes activation and a red arrow denotes inhibition. The legends explain the role of different bounding boxes.

**Fig. 2:**
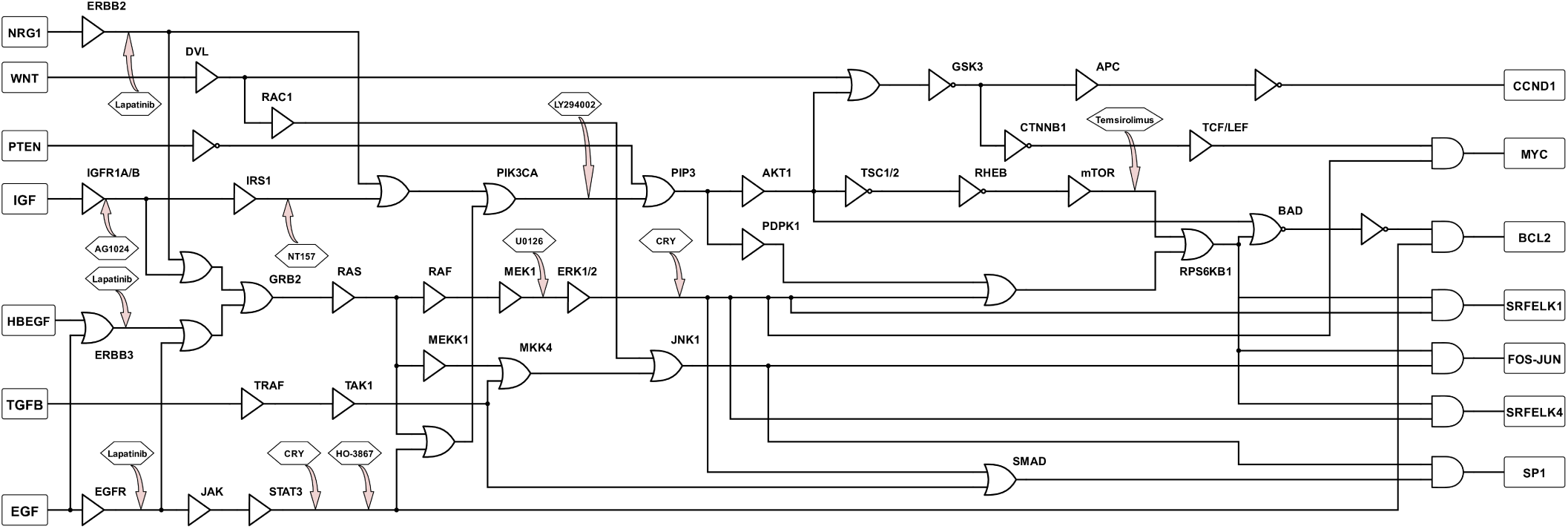
Boolean equivalent of colorectal cancer pathway. The interactions among different genes is modeled using appropriate boolean logic functions.

We now discuss our model simulations with theoretical results and the experimental validation on human colorectal cancer cell lines.

## 4 Results

### Theoretical Results

We computed the ratio of probabilities (ROP) for all the genes in our CRC pathway using equations 1 and 2 and tabulated them in the supplementary materials (see Additional File 1). We further overlaid these ROP values on our boolean network using a color-coded scheme for a better insight as shown in Fig. 3. We assigned a green color for an ROP *<* 1 and a red color for an ROP *>* 1. A gene with green coding (ROP *<* 1) is down-regulated in the tumor samples as compared to the healthy samples, whereas, a gene with red coding (ROP *>* 1) is up-regulated in the tumor samples as compared to the healthy samples. From the figure, it is apparent that the JAK/STAT3, the PI3K/AKT/mTOR, and the RAS/RAF/ERK pathways are highly up-regulated and are possibly the main reason for an out of control cell proliferation in colorectal cancer.

**Fig. 3:**
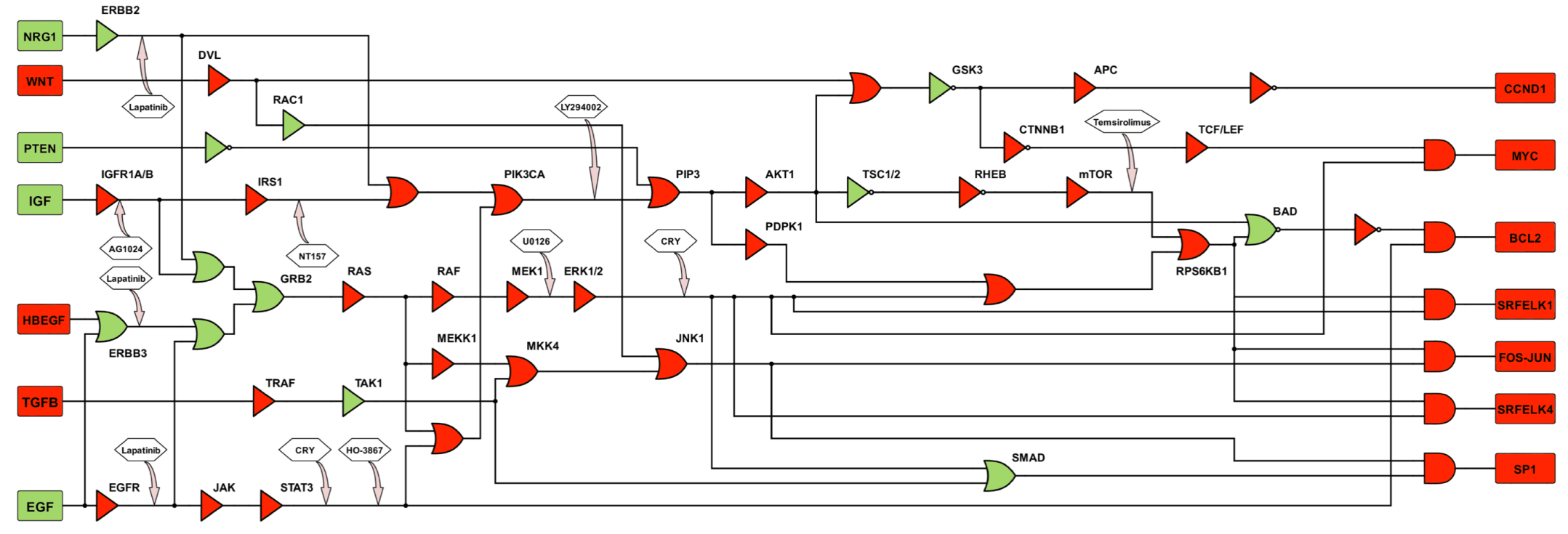
Ratio of probabilities overlaid on the boolean pathway. Here, a green color denotes an ROP *<* 1 and a red color denotes an ROP *>* 1.

After establishing the key genes and pathways that can most likely mutate and override programmed cell death, we are now interested in deducing the drug combination(s) that can negate this oncogenic effect. We simulated our boolean network with three possible mutations occurring simultaneously and obtained the drug combination(s) that efficiently drove the mutated network towards cell death.

Referring to the boolean model in figure 2, we have seven inputs and seven outputs. The inputs consist of six growth factors (EGF, TGFB, HBEGF, IGF, WNT, NRG1) and a tumor suppressor gene (PTEN). The outputs include crucial genes (CCND1, cMYC, BCL2, SRF-ELK1, FOS-JUN, SRF-ELK4, SP1) responsible for cell proliferation and apoptosis. We also have different therapeutic drugs (AG1024, U0126, NT157, LY294002, HO-3867, Lapatinib, Temsirolimus, Cryptotanshinone) intervening at the appropriate targets.

We can mathematically represent these inputs, outputs, and drugs as row vectors. A zero represents an inactive gene/drug in the corresponding location, whereas, a one represents an active gene/drug. Hence, the input, output, and drug vectors are given by:

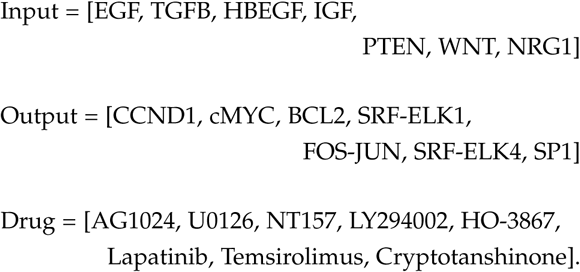

From the boolean network in figure 2, in a healthy cell with the growth factors inactive and the tumor suppressor active (Input = [0000100]), all the resulting output genes are inactive (Output = [0000000]) and this represents an absence of cell proliferation and a non-suppression of apoptosis. However, for the same input, the boolean network with mutations (faults) will generate a non-zero output vector. Our goal is to drive this non-zero output vector closer to the zero vector with the help of therapeutic drugs. Biologically, this translates to using drug intervention to drive a mutated pathway close to cell death. In order to do that, we computed a measure called Size Difference (SD). Let 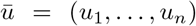 and 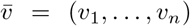 be two binary vectors. Then, we can count the number of matches and mismatches at each bit location and compute SD using equation 3, where *n*_01_ counts the number of vector components with *u*_*i*_ = 0 and *v*_*i*_ = 1 simultaneously. Similarly, we can compute the other components *n*_10_, *n*_00_, and *n*_11_.

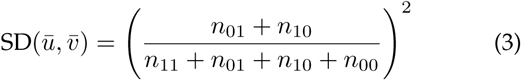

Basically, SD quantifies how different two vectors are and we are interested in computing how different a non-zero output vector is from the ideal output. A higher size difference value correlates with a higher deviation from the ideal output and corresponds to greater cell proliferation and/or curtailed apoptosis. Similarly, a smaller value correlates with a lower deviation from the ideal output and corresponds to decreased cell proliferation and/or increased apoptosis.

We simulated our boolean network for every combination of at most three faults occurring simultaneously and at most three drugs intervening simultaneously. Since there are 33 possible genes that could mutate and 8 drugs in our network, we considered a total of ^33^*C*_1_ + ^33^*C*_2_ + ^33^*C*_3_ = 6017 combinations of mutated networks and a total of ^8^*C*_1_ + ^8^*C*_2_ + ^8^*C*_3_ = 92 combinations of drugs. Hence, we constructed a 6017 *×* 92 matrix of SD values. Finally, we arrived at the best drug combination by choosing the drugs that had the lowest average of size differences across all possible faults. We normalized these average SD values with no therapy (untreated cell line) as the reference and tabulated them in supplementary materials (see Additional File 2), and as representative samples, we present the most notable drugs and their average SD values in table 2.

**TABLE 2:** Drug combinations and their average SD values.

| Drug combinations | Average SD value |
| --- | --- |
| Untreated | 1.000 |
| AG1024 | 0.914 |
| NT157 | 0.990 |
| AG1024 + NT157 | 0.904 |
| U0126 | 0.433 |
| NT157 + U0126 | 0.424 |
| H03867 | 0.875 |
| Lapatinib | 0.680 |
| Temsirolimus | 0.994 |
| Temsirolimus + AG1024 | 0.908 |
| LY294002 | 0.779 |
| LY294002 + H03867 | 0.700 |
| Cryptotanshinone | 0.273 |
| Cryptotanshinone + H03867 + NT157 | 0.263 |
| Cryptotanshinone + AG1024 | 0.251 |
| Cryptotanshinone + Temsirolimus + Lapatinib | 0.193 |
| Cryptotanshinone + Lapatinib | 0.202 |
| Cryptotanshinone + Lapatinib + AG1024 | 0.175 |
| Cryptotanshinone + LY294002 + U0126 | 0.106 |
| Cryptotanshinone + LY294002 + Temsirolimus | 0.087 |

From the table, we see that the drugs AG1024, NT157, HO3867, Lapatinib, U0126, Temsirolimus, and LY294002 have relatively high average SD values suggesting minimal impact and low efficacy. However, Cryptotanshinone, by itself or in combination with other drugs, produced relatively low SD values promising enhanced cell death in CRC cells by targeting the key oncogenic pathways. The detailed codes of this simulation and its implementation are available online at https://github.com/hashwanthvv/colorectal.

### Experimental Results

We corroborated our theoretical results obtained above by carrying out experiments on HT29 and HCT116 human colorectal cancer cells exposed to different drug treatments. We used a high-content fluorescent protein reporter imaging approach and detected cell death in the CRC cells. Using a two-step data processing technique introduced by Hua et al. in 2012 [43], we obtained the cell processing dynamics and compressed this data into expression profiles and plotted them for better elucidation.

The drugs and their combinations we used in our experiments are, AG1024 (20 *µ*M), NT157 (10 *µ*M), U0126 (10 *µ*M), HO-3867 (10 *µ*M), Lapatinib (5 *µ*M), Temsirolimus (10 *µ*M), LY294002 (10 *µ*M), and Cryptotanshinone (20 *µ*M). The drug dosage levels of Lapatinib and Temsirolimus are those of human medical usage, and the dosages for the rest of the drugs are at levels similar to the tests of their utilities on human cell lines.

In figure 4, we plotted the Apoptosis fraction (amount of cell killing) produced in HT29 colorectal cancer cells under the influence of different drug combinations over time. The black line represents the untreated cell line which serves as a reference. Since our theoretical results predicted the effectiveness of Cryptotanshinone, we used it in each of the drug combinations and from the plots, it is evident that impressive cell death occurs within 9-10 hours in each combination containing Cryptotanshinone.

**Fig. 4:**
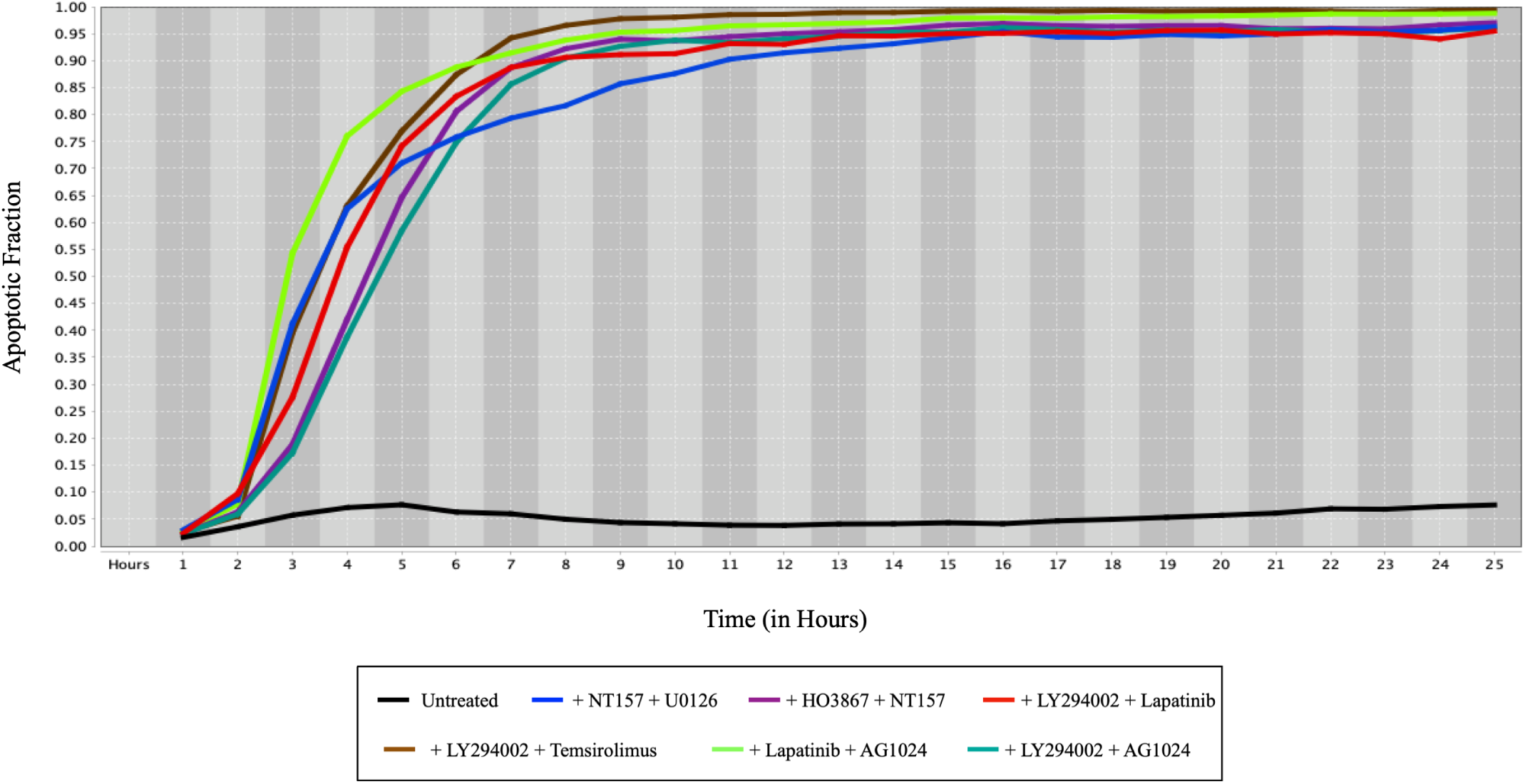
Apoptosis fraction versus time (in hours) for different drug combinations in HT29 human colorectal cells. Cryptotanshinone (CRY) has been used in each of the drug combinations. The drug combinations in the legend from left to right are Untreated cell line, Cryptotanshinone+NT157+U0126, Cryptotanshinone+HO3867+NT157, Cryptotanshinone+LY294002+Lapatinib, Cryptotanshinone+LY294002+Temsirolimus, Cryptotanshinone+Lapatinib+AG1024, and Cryptotanshinone+LY294002+AG1024.

To further validate our argument, we conducted experiments using drug combinations with and without Cryptotanshinone on the HCT116 cell line. We then plotted the results in figure 5, and it is apparent that the drugs, NT157, Temsirolimus, AG1024, are quite futile by themselves, but when complementing those drugs with Cryptotanshinone, we notice an outstanding efficacy of inducing cell death. These results further support our claim that Cryptotanshinone significantly enhances cell death by intervening with the crucial oncogenic genes in colorectal cancer.

**Fig. 5:**
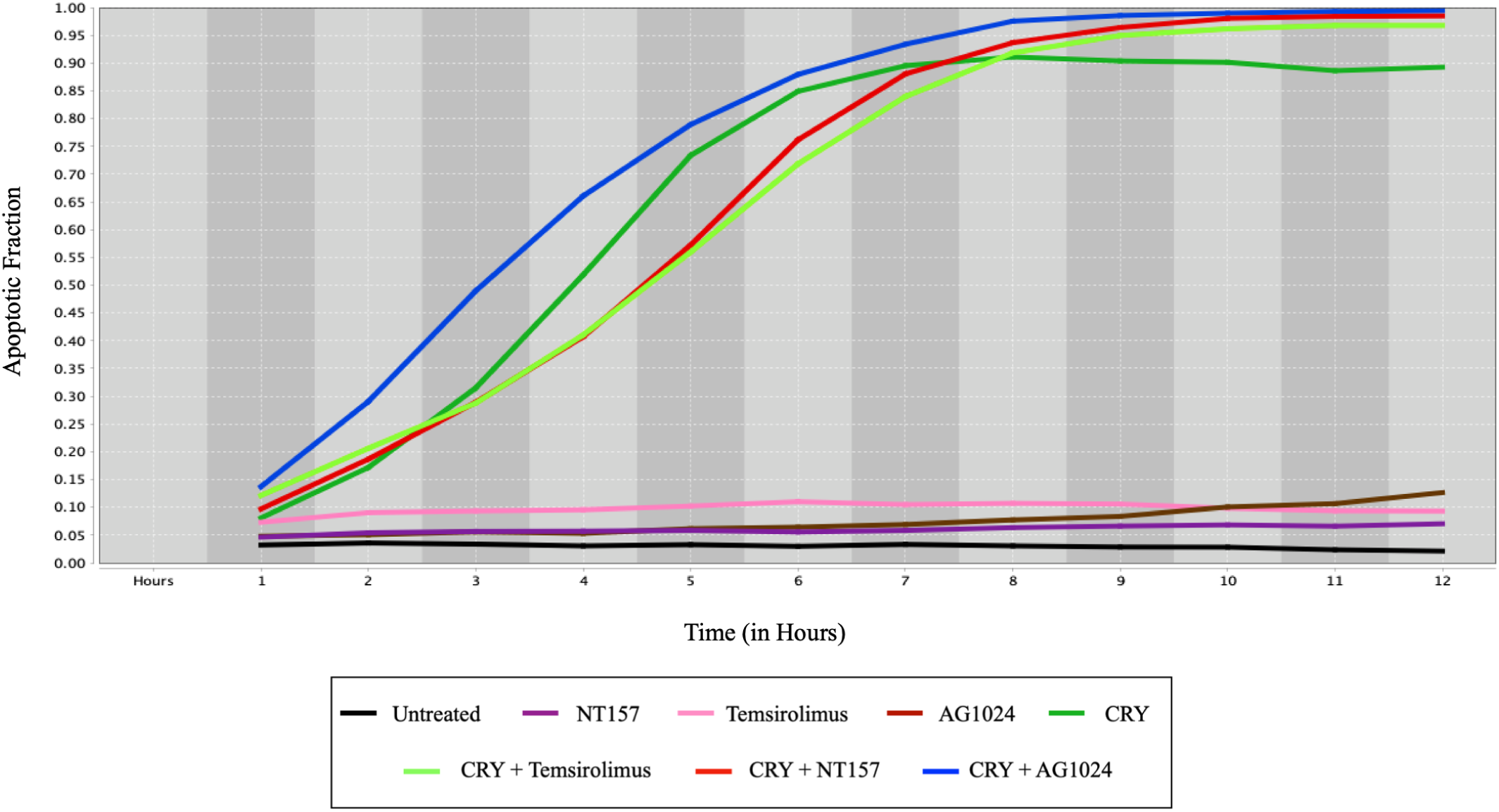
Apoptosis fraction versus time (in hours) for different drug combinations in HCT116 human colorectal cells. The drug combinations in the legend from left to right are Untreated cell line, NT157, Temsirolimus, AG1024, Cryptotanshinone (CRY), Cryptotanshinone+Temsirolimus, Cryptotanshinone+NT157, and Cryptotanshinone+AG1024.

## 5 Discussion

The overall incidence of CRC continues to be on the rise and the 5-year relative survival rate remains lower than 50% in low-income countries. It is predicted that by 2030, the global burden of colorectal cancer will rise by 60% in developing countries [44]. Although surgery and chemotherapy remain the predominant conventional treatments, it is vital to understand the fundamental genes and pathways in CRC to develop more effective treatment approaches.

In this paper, we presented a mathematical framework to model the colorectal cancer pathway and infer the underlying mutations responsible for the tumor growth and progression. Using a public gene expression dataset, we compared the activity of genes in healthy samples versus tumor samples. Our analysis exposed the underlying mutations in the JAK/STAT, the MAPK, and the PI3K pathways as the primary drivers for the uncontrolled cell proliferation. In order to target these pathways, we further modeled the pathway into a boolean technique and deduced the drug combinations which can force the tumorigenic network into cell death. Our simulations showed that Cryptotanshinone, either by itself or in combination, targets the key mutations and halts cell development. We now discuss the biological significance of our results.

Colorectal cancer is a heterogeneous disease defined by different malignancies in RTKs or downstream genes of RTK-activated intracellular pathways. Anti-EGFR agents are often futile because of RAS mutations that occur downstream of EGFR, and these mutations along with other abnormal pathways contribute to CRC manifestation and growth. RAS protein operates in both the mitogen-activated protein kinases (MAPK) and the phosphoinositide-3 kinase (PI3K) pathways, which are crucial in overseeing important cellular functions such as cell differentiation, survival, and growth in healthy cells [24]. In molecular terms, mutations in RAS codons could require conformational changes so that the GTP molecules are not hydrolysed and they maintain RAS constantly in its active state, thus amplifying MAPK signaling and oncogenic effects [23]. RAS can also bind to p110, the catalytic subunit of the PI3Ks, and activate the PI3K pathway. Mutations in RAS makes it difficult to bind to p110 and this forces PIK3CA to undergo a mutation and encode a truncated p110 which then binds to RAS.

The Janus family of tyrosine kinases (JAK)/signal transducer and activator of transcription (STAT) is another critical pathway required for healthy functioning of a cell. Constant activation of STAT3 and/or JAK2 are associated with cell proliferation in several cancers including breast, pancreatic, and lung [26]. This overwhelming data provides novel evidence that the JAK/STAT3 pathway can be a new potential therapeutic target in CRC.

These above significant pathways interact with one another and play significant roles in cellular processes. However, these pathways can get mutated in multiple ways: (1) by upregulation of MAPK/PI3K/STAT genes; (2) by indirect upregulation of activator genes or (3) by down-regulation of inhibitor genes. With myriad possibilities of gene mutations, it is evident that cancers do not respond to general therapy because of downstream malignancies or resistance to therapies like anti-EGFR drugs in CRC [45]. As a result of these diverse activated pathways, a combination therapy targeting multiple mutations seems a promising choice. However, a comprehensive search of the best drug combination is both expensive and tedious. Hence, we used a theoretical model to reduce the search space of drug combinations that guarantee cell death. Our simulations using a boolean model indicated that Cryptotanshinone, a naturally occurring extract from Salvia miltiorrhiza Bunge, halted cell proliferation by inhibiting the crucial pathways.

Cryptotanshinone (CRY) is derived from a traditional Chinese herb and has been shown to have substantial pharmacological effects, including in acute ischemic stroke, chronic hepatitis, and Alzheimer’s disease [46]. CRY has also had significant efficacy in achieving cell death in several human cancer cells including lung, pancreatic, breast, prostate, colorectal, and gliomas [47]. This effectiveness of CRY against multiple cancers is mainly because of its ability to suppress JAK/STAT3 pathway. Cryptotanshinone blocks STAT3’s dimerization preventing it from regulating its downstream proteins [41]. CRY also induces potent growth inhibition by decreasing cyclin A1 and increasing cyclin D1 protein levels [48]. Therefore, our results provide supporting evidence of Cryptotanshinone and its use as a potential therapeutic agent in cancer.

The literature and the above discussion substantiate our claims that Cryptotanshinone by itself and in combination with other therapeutic agents is a promising drug in the treatment of colorectal cancer. We conclude that our theoretical and experimental findings prove the efficacy of Cryptotanshinone and believe that these models can make way for the advancement of new and better techniques for pathway modeling and ranking drug efficacies in biological networks.

## Acknowledgment

This work was supported in part by the US National Science Foundation under Grants ECCS-1609236 and ECCS-1917166 and in part by the TEES-AgriLife Center for Bioinformatics and Genomic Systems Engineering (CBGSE) startup funds. The statements made herein are solely the responsibility of the authors.

